# The metabolic pace of life histories across fishes

**DOI:** 10.1101/2020.11.16.385559

**Authors:** Serena Wong, Jennifer S. Bigman, Nicholas K. Dulvy

## Abstract

All life acquires energy through metabolic processes and that energy is subsequently allocated to life-sustaining functions such as survival, growth, and reproduction. Thus, it has long been assumed that metabolic rate is related to the life history of an organism. Indeed, metabolic rate is commonly believed to set the pace of life by determining where an organism is situated along a fast-slow life history continuum. However, empirical evidence of a relationship between metabolic rate and life histories is lacking, especially for ectothermic organisms. Here, we ask whether three life history traits – maximum body mass, generation length, and growth performance – explain variation in resting metabolic rate (RMR) across fishes. We found that growth performance, which accounts for the trade-off between growth rate and maximum body size, explained variation in RMR, yet maximum body mass and generation length did not. Our results suggest that measures of life history that encompass trade-offs between life history traits, rather than traits in isolation, explain variation in RMR across fishes. Ultimately, understanding the relationship between metabolic rate and life history is crucial to metabolic ecology and has the potential to improve prediction of the ecological risk of data-poor species.

## Introduction

Metabolism is the process by which all living organisms turn external resources into available energy and, in turn, allocate this energy among competing life history processes, such as survival, growth, and reproduction [1–3]. One theory, the Metabolic Theory of Ecology (MTE), proposes a mechanistic basis for understanding why metabolic rate scales with body mass with an exponent of three quarters (0.75) [4]. From this, the MTE derives quarter-power scaling relationships for numerous ecological phenomena including life history traits [4,5]. The MTE thus assumes that metabolic rate can be used as a predictive tool to understand traditionally difficult-to-measure ecological phenomena, based upon the similarity of scaling exponents. However, predictions of many higher-order ecological phenomena are based on the underlying assumption that metabolic rate underpins life history. Surprisingly, there are few empirical tests of whether life histories are directly related to metabolic rate, particularly for ectotherms [6–8]. If this putative relationship between metabolic rate and life histories exists, the idea that there is an organismal physiological basis underlying conservation and global-change-related phenomena, such as overfishing, climate change responses, and extinction risk, may prove to be a reality [2,9–11]. Thus, exploring the connections between metabolic rate and life histories may increase our understanding of the diversity of life histories and offer simple, trait-based approaches to support the development of ecological risk analyses [2,12].

Life history traits are optimized through natural selection to maximize fitness [2,3,9]. Trade-offs among life history traits arise as individuals have finite resources to allocate to the competing processes related to survival, growth, and reproduction [2,13,14]. For example, there is a trade-off between maximum size and growth rate whereby fishes either grow fast to a small size or grow slower to a larger size [15,16]. In turn, these trade-offs and the environment provide the framework for the evolution of life history traits [2,9]. Specifically, in response to selection imposed by a particular environment, suites of life history traits commonly co-evolve, clustering together along a fast-slow axis, with organisms that grow slower, mature later, live longer, and have a larger maximum body size on the ‘slow’ end of the continuum, and organisms with the opposite suite of traits on the fast end [2,13,17,18]. Thus, life history traits can characterize an organism’s pace of life, as they describe where an organism is situated along this fast-slow continuum of life history [13]. The MTE predicts that metabolic rate sets this pace of life, thus determining where organisms sit on a fast-slow continuum. Under this theory, organisms with a higher metabolic rate will sit towards the ‘faster’ end of the life history continuum, since allocation of resources to growth and reproduction is powered by a faster metabolism [4]. Yet, the relationships between metabolic rate and life histories have rarely been examined across species, and when they have, it has yielded conflicting results. For endotherms (birds and mammals), it is still unclear whether age-related life history traits such as age at first reproduction and maximum age are related to metabolic rate, even after controlling for body mass and evolutionary history [6,8,19]. Conflicting results have also been found in studies of growth rate. While growth rate has been found to be a strong, positive predictor of resting metabolic rate across vertebrates and has also been found to positively correlate with metabolic rate in nestling songbirds, in other studies of birds and mammals no relationship has been found between growth rate and metabolic rate [6,7,20,21]. Furthermore, we know little about relationships between metabolic rate and life histories for ectotherms. Fishes present a unique opportunity to examine this relationship in ectotherms, as they are the most speciose group of ectotherms, constitute one of the most taxonomically and metabolically diverse radiations of vertebrates, and exhibit a wide range of life histories [22–24]. Thus, examining whether metabolic rate and life history traits are related across fishes allows us to test a fundamental premise of metabolic ecology in ectotherms.

Here, we ask whether life history traits explain variation in metabolic rate across fishes, after accounting for shared evolutionary history and the effects of body mass and temperature. Specifically, we examined whether three life history traits – maximum body mass, generation length, and growth performance – were related to resting metabolic rate (RMR) across 104 fish species using a phylogenetic generalized least squares regression framework [25]. We hypothesized that all three life history traits would explain variation in RMR, but that growth performance would explain the most variation in RMR because it encapsulates a life history trade-off (between growth and maximum size), whereas maximum body mass and generation length do not encapsulate trade-offs. Specifically, we predicted that species with a high metabolic rate for their body mass would have the characteristics of a ‘faster’ life history – a smaller maximum body mass, a shorter generation length, and a higher growth performance.

## Materials and methods

### Metabolic rate data collation and selection

Resting metabolic rate (RMR), measurement temperature (i.e., the temperature associated with the metabolic rate measurement), and measurement body mass (i.e., the wet body mass associated with the metabolic rate measurement) were collated from the literature. For our analysis, we only used estimates of RMR from rates of oxygen consumption for post-absorptive, post-larval fishes in which oxygen uptake due to activity was mitigated. Obtaining estimates of RMR only from peer-reviewed studies allowed us to categorize the type of metabolic rate (e.g., RMR) measured in each study with a high degree of confidence and avoid propagating potentially erroneous metabolic rate estimates. We collated raw data (i.e., separate estimates for individuals of the same species) for each of the three metabolic traits, if available, although in most cases only a species’ mean was published. Thus, for our analyses, we averaged raw estimates of RMR and measurement body mass at a given measurement temperature, resulting in a species-specific mean RMR, measurement temperature, and mean measurement body mass. If more than one study reported RMR for the same species, we chose only one study to include in our dataset to avoid biasing our results towards species that were represented by multiple studies, following Killen et al. [22]. To ensure that our choice of which study for a given species to include in our dataset did not affect the results, we conducted all analyses on three separate datasets (the ‘sample size dataset’, the ‘mass dataset’, and ‘the temperature dataset’) resulting from the following inclusion criteria: (1) based on the largest sample size, as presented in the main manuscript, (2) based on the largest average measurement body mass, to approximate maximum body size, and (3) based on which study’s measurement temperature was within the natural temperature range of the species but closest to 20°C to minimize the range of temperatures included in the dataset following Gillooly et al. [26] and Killen et al. [22]. If a study measured RMR at multiple measurement temperatures, we also used selection criteria to determine which RMR data to include. For more detail on data collation and our selection criteria see section 1 of the Supplementary Methods.

### Life history data collation, selection, and aggregation at the species level

To assess if life history traits explain variation in RMR, we collated maximum body mass, generation length, and growth performance collected from peer-reviewed studies and grey literature (hereafter ‘life history study’) using literature searches and FishBase [16,27]. These life history traits were chosen because they are available for many species and are widely used to describe fishes’ life histories [15,16,28]. Maximum body mass was collated from the literature or derived from maximum body length using species-specific length-weight conversions (for more detail, see Supplementary Methods, section 2.1). Generation length and growth performance are both life history traits that are calculated from multiple other life history traits (i.e., they are both ‘composite’ life history traits). Generation length was calculated as *T_mat_* + *T_max_* * *z*, where *T_mat_* is age at maturity, *T_max_* is the maximum age recorded for the species, and *z* is a constant that depends on survivorship and the relative fecundity of young versus old individuals in the population [29,30]. We used a conservative value of *z* = 0.5 that is consistent with IUCN guidelines to account for the truncation of age structure in many fish populations by overfishing (Supplementary Methods, section 2.2) [29,31]. Growth performance is a composite life history trait that allows for the comparison of growth rates across species that differ in maximum size, and thus accounts for the trade-off between growth and maximum size [32,33]. Growth performance is often calculated as phi prime, *ϕ’* = log_10_(*k*) + 2*log_10_(*L_∞_*), where *L_∞_*, is asymptotic length, or the mean body length that individuals in the population would reach if they were to grow indefinitely, and *k* (year^−1^) expresses the rate at which the asymptotic length is approached [32,33]. We also calculated growth performance using another common measure, yet our analysis was largely insensitive to this choice (Supplementary Methods 2.3). Finally, for 28 of the 104 fish species, not all life history traits were available and thus life history trait values from closely related species (here, ‘proxy species’) were used. To ensure that our results were not sensitive to the inclusion of data from proxy species, we reran analyses while excluding all species for which life history trait data from proxy species was used, and compared results (Supplementary Methods, section 2.4).

### Statistical Analyses

We included a phylogenetic random effect in all models to account for phylogenetic non-independence among residuals using phylogenetic generalized least squares (PGLS) as implemented in the *caper* package [25,34]. We constructed a supertree from two sources: (1) the teleost Fish Tree of Life [23], and (2) a molecular phylogeny for chondrichthyans [24] using the R package *phytools* [35]. All statistical analyses were conducted in R v. 3.6.1 [36].

#### 1. Do life history traits explain variation in RMR across fishes?

To test whether life history traits explain variation in RMR across fishes, we parameterized and compared four models – one for each of the three life history traits (i.e., maximum body mass, generation length, or growth performance), and a ‘null model’. The ‘null model’ included only measurement body mass and measurement temperature as explanatory variables. For each life history model, RMR was the response variable, and measurement body mass, measurement temperature, and the respective life history trait were the explanatory variables. For all models, measurement body mass was converted to grams, measurement temperature was converted to inverse temperature, 1/(temperature\**K*), where *K* = Boltzmann’s constant and temperature is in Kelvin following Gillooly et al. [26], and then standardized, and RMR was converted to watts following Grady et al. [21]. All variables, other than inverse measurement temperature and growth performance, were log_10_-transformed for all models. It should be noted that growth performance is already on a log_10_ scale by nature of its calculation. Comparisons of the four candidate models were then made using Akaike information criteria (AICc), which penalizes models for their number of estimated parameters, with smaller AICc values indicating a better model fit [37]. Of the candidate models, the weight of evidence for any given model was measured by its Akaike weight (*w_i_*), the relative likelihood of the model divided by the sum of the likelihoods of all other models. Finally, as generation length and growth performance are composite life history traits, we parameterized four additional models – two with the components of generation length (i.e., age at maturity and maximum age) and two with the components of growth performance (i.e., *k* and *L_∞_*) as explanatory variables – to ensure that no one component of these composite traits was driving the relationship with RMR.

#### 2. What is the relative importance of each life history trait in explaining variation in RMR across fishes?

To assess the relative importance of maximum body mass, generation length, and growth performance in explaining variation in RMR across fishes, we fitted a model that included measurement body mass, measurement temperature, maximum body mass, generation length, and growth performance as explanatory variables (hereafter, ‘global model’). Collinearity between variables was checked using variance inflation factors (VIFs) and all VIFs were less than five [38]. All explanatory variables were centered and scaled by subtracting the mean and dividing by twice the standard deviation (hereafter ‘standardized’) so that effect sizes could be interpreted and compared in terms of units of standard deviations [25,39].

### Results

#### 1. Do life history traits explain variation in RMR across fishes?

Overall, we found that the only life history trait which explained variation in RMR across fishes was growth performance, which encompasses a life history trade-off (Figure 1A). The best overall model (AICc = 16.84, *wi* = 0.989; Table S1) described RMR as a function of measurement body mass, measurement temperature, and growth performance. Growth performance explained variation in RMR even after accounting for the effects of measurement body mass and measurement temperature (Figure 1B), with species with a high metabolic rate for their measurement body mass also having a high growth performance (*β* = 0.24, 95% confidence interval, CI: 0.12 – 0.36; Table S2). On the other hand, the other life histories traits did not explain variation in RMR despite our prediction that species with a high metabolic rate for their measurement body mass would have a smaller maximum body mass and a shorter generation length. Specifically, a null model with only measurement body mass and measurement temperature had similar relative support (AICc = 27.47, *w*_i_ = 0.005) to the models containing either maximum body mass (AICc = 27.80, *w*_i_ = 0.004), or generation length (AICc = 29.58, *w*_i_ = 0.002, Table S1). Thus, maximum body mass (Figure 1C) did not explain variation in RMR after accounting for the effects of measurement body mass and measurement temperature (Figure 1D) and generation length (Figure 1E) did not explain variation in RMR after accounting for the effects of measurement body mass and measurement temperature (Figure 1F). Similarly, none of the component traits – age at maturity, maximum age, *k* or *L_∞_* – used to calculate the composite traits of generation length and growth performance explained variation in RMR on their own, as the 95% CI of their effect sizes crossed zero (Table S2).

**Figure 1.**
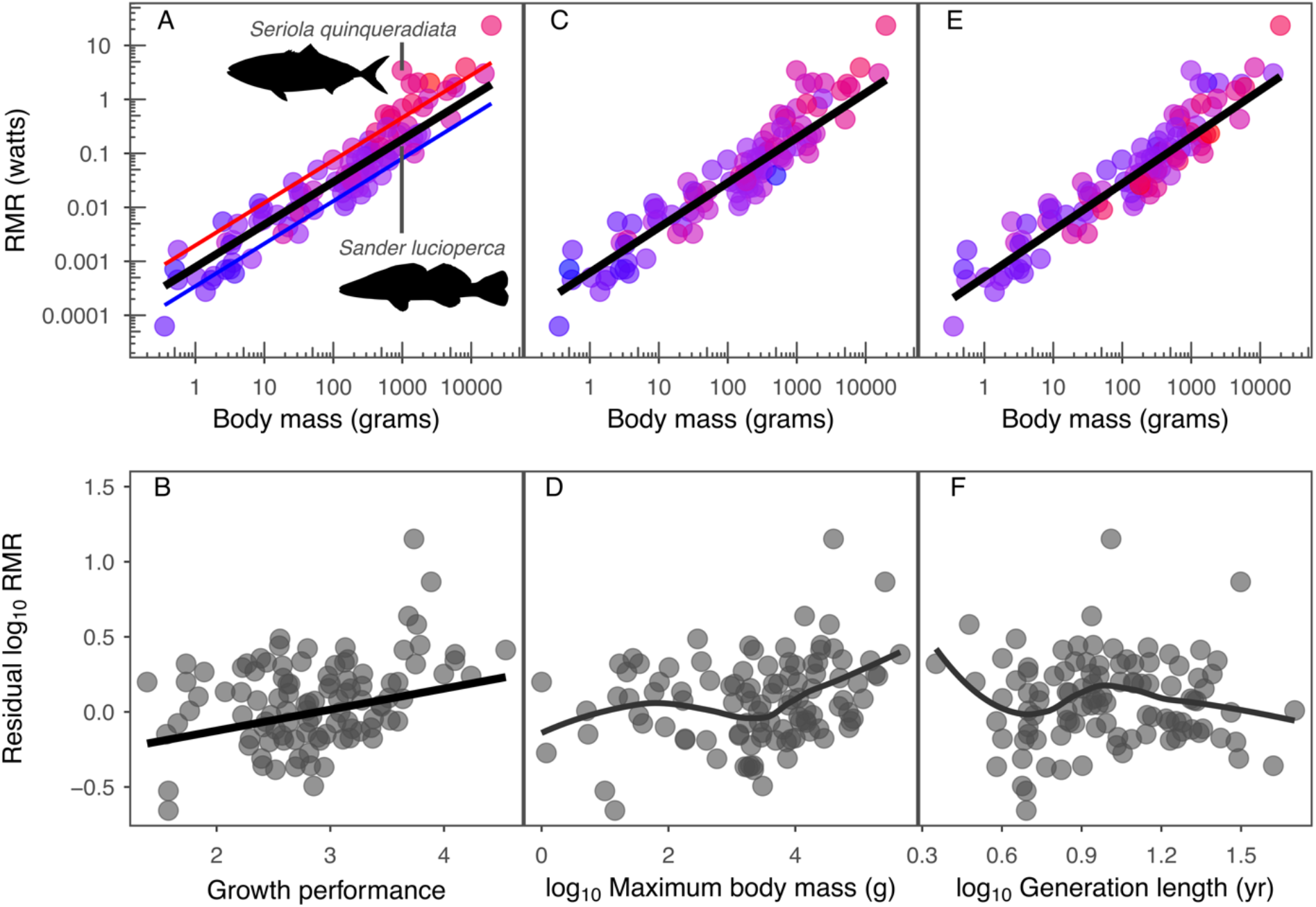
Relationships between resting metabolic rate (RMR), measurement body mass, and life history across fishes. Points are coloured by growth performance (A), maximum body mass (C), and generation length (E), where red denotes larger values and blue denotes smaller values. The black fitted regression line in all three panels is the estimated RMR across the body sizes of all species in the dataset, while incorporating temperature, evolutionary history, and the relevant life history trait (maximum body mass, generation length, or growth performance). Growth performance was the only life history trait to explain variation in RMR, as illustrated by the red and blue lines in (A) which show the estimated RMR for species with high (e.g. Japanese Amberjack, *Seriola quinqueradiata*) and low (e.g. Zander, *Sander lucioperca*) values of growth performance while accounting for temperature and evolutionary history. The bottom row shows residual RMR after accounting for measurement body mass and measurement temperature with linear model (B) and loess (D, F) lines. (Online version in colour.)

#### 2. What is the relative importance of each life history trait in explaining variation in RMR across fishes?

Only growth performance, measurement body mass, and measurement temperature explained variation in RMR, as evidenced by their relative effects in a global model with standardized explanatory variables (Figure S1, [39]). Growth performance had over a four-fold larger effect on RMR compared to maximum body mass, and a 34-fold larger effect on RMR compared to generation length (Figure S1).

#### Sensitivity analyses

Our findings were robust to the three different data inclusion criteria (Supplementary Results, Table S3), an additional measure of growth performance (Tables S4, S5), and the use of traits from related proxy species to in-fill data gaps (Table S6). Finally, the residuals from all models had a phylogenetic signal (lambda, *λ*) of 0.56 or greater, indicating that including a random effect of phylogeny is necessary when examining metabolic rate across species (Table S1).

### Discussion

Our study directly tests whether life history explains variation in RMR across fishes, and our findings help reconcile the conflicting results of previous work relating metabolic rate and life histories across species. We find that the connection between metabolic rate and life histories across fishes only exists when accounting for life history trade-offs, such as the trade-off between growth rate and maximum size, and that neither maximum body mass nor generation length explained variation in RMR after accounting for measurement body mass, measurement temperature, and evolutionary history. First, we compare the relationships among various measures of life history and RMR and discuss these results in the context of life history trade-offs. Second, we consider the utility of this and other studies for explaining broad life history patterns and the implications for metabolic ecology. Finally, we highlight future directions for furthering our understanding of the relationships between metabolic rate and life histories.

We found that of the life history traits examined, only growth performance explained variation in RMR across fishes. We hypothesized that growth performance would explain this variation because it incorporates a trade-off between life history traits (i.e., between maximum size, *L_∞_*, and growth rate, *k*) and thus may better characterize a fishes’ life history strategy [13,16,32,40]. In contrast, when the components of growth performance (*k* and *L_∞_*) were examined in isolation, they did not explain variation in RMR, emphasizing the need to examine composite indices that encompass trade-offs when investigating the relationship between RMR and life history. However, some composite traits may not fully capture life history trade-offs among competing processes. For example, generation length is also a composite measure of life history that combines age at maturity and maximum age, yet it did not explain variation in RMR, likely because it does not capture a life history trade-off. The lack of relationship between RMR and maximum body mass in our study was also notable because maximum body mass is widely used as an indicator of an organism’s position along the fast-slow life history continuum and is often used in assessments of extinction risk in ectothermic species [16,28]. Instead the size-dependency of metabolic rate may be mostly captured by measurement body mass, leaving little variation to be explained by maximum body mass, despite the differences in these two measures. Consequently, empirical tests of the foundations of the MTE should explicitly consider life history trade-offs in fishes, and potentially other ectotherms, rather than individual life history traits in isolation.

Testing the assumption that metabolic rate sets the pace of life histories provides insight into broad life history patterns, such as the temperature-size rule, and is a first step before using the MTE in its intended predictive capacity. Like metabolic rate, life history traits such as growth rate and maximum size are also temperature-dependent and there is a large body of literature connecting environmental temperature to growth and body size [41,42]. This phenomenon, where individuals grow faster but attain a smaller body size at higher temperatures both in the wild (i.e., latitudinal gradients in body size and growth) and in the laboratory, has come to be known as the temperature-size rule [41,42]. While the mechanistic basis of this phenomenon remains unresolved, hypotheses that connect metabolic processes to life history patterns, such as the oxygen limitation hypothesis, have been proposed, at least for aquatic ectotherms [32,42]. Our results underscore the links between metabolic rate and growth performance and suggest that oxygen consumption may play a role in the temperature-size rule. Additionally, a clearer understanding of whether life history explains variation in metabolic rate across taxa is necessary before the MTE can be reliably used as a predictive model of life histories. If future studies find that life histories explain variation in metabolic rate for both endotherms and ectotherms, we will then be set with the challenge of determining whether (1) metabolic rate does indeed dictate and drive life history, (2) life history drives metabolic rate, (3) metabolic rate and life history are co-adjusted with each other, affecting each other in a reciprocal manner, or (4) both life history and metabolism are indirectly related to additional factors [43]. These studies will not only require correlative approaches as executed here, but selection and common-garden experiments to uncover mechanistic drivers.

Other measures of metabolic rate, life history trade-offs, and statistical approaches may help clarify the relationship between metabolic rate and life history in the future. First, RMR, while the most commonly reported measure of metabolic rate, only reflects energy use and availability at rest, and does not describe the scope for processes such as activity, growth, or reproduction [1]. Field metabolic rate, for example, is likely a more accurate measure of day-to-day energy expenditure than RMR and thus could be more closely linked to life history strategy than RMR [1]. Second, while our results indicate that a measure of life history that accounts for a trade-off explains variation in RMR, there are other life history trade-offs, popularised as Beverton’s dimensionless ratios or Charnov’s life history invariants [44,45]. For example, natural mortality rate (*M*) has been found to be positively related to *k* from the von Bertalanffy growth function and negatively related to age at maturity, so testing whether invariants such as *M*/*k* and *T_mat_*M* also explain variation in metabolic rate may be a fruitful avenue for future research, especially in taxa for which reliable estimates of mortality rate are available (e.g., phytoplankton or birds, [44,45]). Third, new statistical approaches that explicitly account for trade-offs and correlations between life history traits may help us reconstruct life history strategies for species and populations that are data-poor by estimating difficult-to-measure life history traits, such as fecundity [46,47].

Environmental and ecological factors such as activity levels, predation risk, food availability, and environmental temperature may obscure relationships between metabolic rate and life history traits, particularly in ectotherms, and this could also be considered in the future [22,42,48]. Fish species with a high metabolic rate for their body mass have a high growth performance, but they may also have high activity levels [22,32]. For example, Japanese Amberjack (*Seriola quinqueradiata*) had a higher RMR than Zander (*Sander lucioperca*), though metabolic rates of both species were measured on individuals of similar body masses and at similar measurement temperatures (Figure 1A). This difference in RMR may be because Japanese Amberjack had a higher growth performance than Zander, but activity level may also play a role. Metabolic rate is commonly used as a proxy for activity level, confounding studies of the relationship between metabolic rate and activity level [49]. Thus, future studies should investigate the interrelationships between activity level, metabolic rate, and life history by using morphological proxies of activity such as the caudal fin aspect ratio (= [*height of the caudal fin*]^2^/[*surface area of the fin*]) [22,48]. For example, the caudal fin morphology of the Japanese Amberjack is strongly lunate, suggesting that this species is more active compared to the Zander with its rounded tail (Figure 1A). Additionally, predation risk, environmental stability, and food availability, while sometimes experimentally tractable, are difficult to tease apart, let alone account for in macroecological analyses, despite likely influencing both metabolic rate and life history. However, if realistic approximations of predation risk can be attained, dynamic state variable models may provide an avenue for future investigation by featuring the trade-offs associated with life history and factors such as predation risk and food availability within a dynamic behavioural context to determine fitness [12]. Finally, while measurement temperature greatly affects metabolic rate, environmental temperature may have an evolutionary effect on both metabolic rate and life histories through thermal constraints on production or thermal effects on survival as illustrated by broad patterns such as the temperature-size rule [26,41]. In conclusion, our analyses show that growth performance, but not maximum body mass or generation length, explains variation in RMR across a diverse set of 104 fish species. To our knowledge, this is the most comprehensive study to-date that tests whether empirical measures of life history explain variation in metabolic rate across fishes. Our findings revealed that a measure of life history that incorporates a trade-off between life history traits is strongly associated with RMR and therefore provides some support for the assumption that metabolic rate sets the pace of life across species. Insight into the links between physiology and life histories has the potential to inform ecological risk assessments, particularly for data-poor species, because life histories are closely related to risk of overfishing and extinction risk [15,18,28].

## Authors’ contributions

SW, JSB, and NKD conceived of the study; SW collected the data and performed analyses and visualizations; SW wrote the first draft of the manuscript with input from all authors; JSB and NKD advised on the project and NKD supervised the project. All authors gave final approval for publication and agree to be held accountable for the work performed therein.

## Acknowledgements

The authors thank Dr. John Reynolds and the Dulvy lab at SFU for feedback on the study.

## Media Summary

Metabolic rate is believed to govern the rate that energy is allocated to growth, reproduction, and survival and is sometimes used to predict life history traits like growth rate and lifespan. However, we know little about these complex connections. In the most comprehensive study of these connections across fishes, we found that the only measure of life history that explained variation in metabolic rate incorporated a trade-off between traits. As overfishing and climate change loom, this new understanding may prove useful for prioritizing conservation but shows that the predication of life histories is not as straightforward as initially assumed.

## Supplementary Material

See attached document.

