## Supplementary Material for "The metabolic pace of life histories across fishes"

### Supplementary Methods

#### *1. Metabolic rate data collation and selection*

In most studies, RMR was reported as having been calculated by either extrapolating values of oxygen consumption at varying activity levels to no activity, or by measuring oxygen consumption during periods of quiescence after acclimation in the respiratory chamber. When a study reported metabolic rates for the same species at multiple temperatures, only metabolic rate measured at one temperature was included in our dataset. To choose which metabolic rate and measurement temperature to include, we used multiple inclusion criteria: (1) selecting RMR data based upon whichever measurement temperature had the largest sample size, (2) selecting RMR data based upon whichever measurement temperature had the largest average measurement body mass, and (3) selecting RMR data based upon whichever measurement temperature was closest to 20˚C, following Gillooly et al. [1] and Killen et al. [2]. In cases where metabolic rate was measured multiple times for the same individual at the same measurement temperature, the measurement with either the largest measurement body mass or, measurement body mass being equal, the lowest RMR estimate, was used. Occasionally, measurement body mass or measurement temperature were reported as ranges, in which case the midpoint was used.

*2. Life history data collation, selection, and aggregation at the species level*

*2.1 Maximum body mass*

To obtain maximum body mass for each species, we extracted the maximum observed body size (i.e., maximum body length or maximum body mass) from each life history study and from FishBase [3]. If a range of values was given, the maximum of the range was used as it is the largest observed measurement. To obtain a species-specific value of maximum body mass, we preferentially chose the largest value of maximum body mass provided from published papers, and if that was not available, then values from FishBase were used [3,4]. If values of maximum body mass from neither source were available, values of maximum body length were converted to maximum body mass using species-specific length weight regressions. For this, a length-weight regression equation from the same study that estimated maximum body length was used if it was available. If not, species-specific length-weight regression equations were obtained from FishBase [3]. As Fishbase often reported multiple length weight coefficients for each species, we took an average of these for use in converting length to weight. Each species-specific observation was documented in FishBase as having been estimated using a specific length type (e.g. fork length, standard length, etc.) from samples of all male, all female, mixed sex, or 'unsexed' individuals. Observations that were documented as either mixed sex or unsexed were all treated as mixed sex as it was not possible to know the sex composition of these samples. Mean length-weight coefficients were calculated from 'group-specific' (i.e., male, female, or mixed) data and then used to convert length to weight following the formula $W = aL^{b}$, where W is body mass, L is body length, $a$ is the intercept, and $b$ is the allometric slope.

*2.2 Generation length*

To estimate generation length for each species, we extracted its components, maximum age and age at maturity, from each life history study and from FishBase [3]. We extracted the maximum observed age (empirical longevity, in years) from all life history studies in which age was estimated, as well as the theoretical longevity based on the von Bertalanffy growth function, if reported. If a range of values was given, the maximum of the range was used [4]. We compared the empirical ($T_{max}$) and calculated longevity ($T_{\infty}$) estimates to evaluate their interchangeability and check for any potential errors following Juan-Jordá et al. [4], where longevity was calculated using Taylor's relationship, $T_{\infty} = 3/k$ [5]. To estimate a species-specific value of maximum age, we preferentially chose the maximum measured value provided from published papers, if that was not available, then values from FishBase were used [3,4]. If values from neither source were available, then theoretical values of maximum age provided in peer-reviewed life history studies were used. When extracting age at maturity, we did not differentiate between studies that estimated age at maturity as age at which 50% of the sampled individuals have matured and those that reported age at first maturity, following previous work [4]. If a range of age at maturity values was given, the midpoint of the range was used [2]. To estimate a species-specific value of age at maturity, we used a simple arithmetic mean of values provided from published papers. If no values from papers were available, values from FishBase were used [3]. We preferentially used estimates of generation length and its components for females whenever maximum age and age at maturity were reported separately for sexes [4].

*2.3 Growth performance*

To estimate growth performance for each species, we mined out the $L_\infty$ and *k* parameters from studies that estimated growth using the three-parameter formulation of the von Bertalanffy growth function [6]. We compared the maximum observed length ($L_{max}$) and the theoretical maximum length (i.e., asymptotic length, $L_\infty$) of each species to evaluate their interchangeability and check for any potential errors following Juan-Jordá et al. [4]. We calculated growth performance using $L_\infty$ and *k* individually for each study and then attained a single value for each species by calculating a simple arithmetic mean (giving equal weight to all the studies; [4,7]). In order to collate estimates of $L_\infty$ across studies, we converted all $L_\infty$ estimates to total lengths (TL), as other length measurements are difficult or impossible to measure for some species. Disc width (DW, the maximum width across the body) was used instead of TL for ray-like chondrichthyan fishes, as it is the standard measurement of body length for those species and estimates of TL are prone to error [8]. We converted other length types into cm TL (or DW for ray-like chondrichthyans) using published length-length regression equations following the same protocol as length-weight regression equations (see Supplementary Methods section 2.1). If the life history study provided a length-length regression equation then that was used, otherwise, mean species-specific length-length regression coefficients for each group (i.e., male, female, or mixed) and 'known length' type (e.g. fork length, standard length, etc.) were calculated from FishBase [3]. 'Known length' was converted to TL (or DW for ray-like chondrichthyans) following the formula TL = $a + b$*L, where L is the known body length (e.g. measured as fork length, standard length, etc.), *a* is the intercept of the regression, and *b* is the slope.

We evaluated the reliability of the von Bertalanffy growth parameters of each of the species using two criteria. First, for each life history study that estimated von Bertalanffy growth parameters, we estimated the variability in the ratio between the maximum observed length ($L_{max}$) and asymptotic length ($L_\infty$). We eliminated those studies with ratios which fell more than three standard deviations away from the mean ratio across all studies within each species [4]. Second, we examined the variability of the growth performance parameter ($\phi$') calculated from each study across all studies and with species pooled. The $\phi$' values for a given species or taxonomically related group of species should be normally distributed around the mean $\phi$' of the taxonomic unit, and values further away from the mean of the distribution must be interpreted with increasing caution [4,9,10]. We standardized the $\phi$' values of each study by dividing each by the mean of $\phi$' within each species. Von Bertalanffy growth equations for which the standardized $\phi$' value was greater than three standard deviations away from the mean standardized $\phi$' value for all studies and species were then removed. A cut-off of three standard deviations was chosen based on previous work [4] as well as the histogram of the standardized $\phi$' (all data pooled).

Although we chose phi prime, $\phi$', as a measure of growth performance because it is widely used and thus could facilitate comparisons, it has been suggested that growth performance indices based upon asymptotic weight, $W_\infty$, rather than asymptotic length, $L_\infty$, may be better when comparing fishes that differ in body shape [10–12]. Thus, we re-ran analyses using a measure of growth performance based upon asymptotic weight (phi, $\phi$) in order to compare results for the two measures of growth performance. Phi was calculated using the equation $\phi = \log_{10}(k) + 2/3*\log_{10}(W_\infty)$ [9]. $W_\infty$ was calculated from $L_\infty$ using length-weight regression equations (see

Supplementary Methods section 2.1). As length-weight regression equations were not available for all species and groups (i.e., male, female, or mixed sex), the relationship between growth performance and RMR was investigated using a subset of data for which $\phi$ could be calculated. Additionally, species-specific, but not group-specific, length-weight regression equations were used to estimate $W_\infty$, and thus $\phi$, for a larger subset of species in a separate analysis. Thus, we tested whether weight-based growth performance explained variation in RMR using two datasets – one where $\phi$ was calculated from species-specific and group-specific length-weight regression equations (n = 44) and one $\phi$ was calculated from species-specific but not group-specific length-weight regression equations (n = 84).

#### *2.4. Proxy species*

For the 28 species that were missing at least one life history trait, values from closely related species (i.e., 'proxy species') were mined out from life history studies of species within the same genus as the species with the missing trait value. Life history trait values from each life history study were then aggregated following the methods outlined in Supplementary Methods section 2.

### **Supplementary Results**

Our results were similar regardless of the method of data selection used (Figure S1, Table S1-3), were generally robust to the measure of growth performance used (Table S4, S5), and were robust the inclusion of life history data from proxy species (Table S6). When growth performance was characterized using a weight-based metric ($\phi$), rather than a length-based metric ($\phi$'), results seemed to depend upon sample size. When $\phi$ was calculated using estimates of $W_{\infty}$ from group- and species-specific length-weight regressions, our sample size was more than halved (n = 104 vs. n = 44). Thus, the candidate models all had similar support (all ΔAICcs were less than two and models had similar Akaike weights) and $\phi$ was not correlated with RMR (Table S4, S5). However, when species-specific, but not group-specific, length-weight regressions were used, we were able to obtain estimates of $\phi$ for a larger dataset (n = 84) and our results matched analyses using $\phi'$. When models including either $\phi$ or $\phi$' were compared with AICc, they both had similar relative support and were within two AICc of each other (Table S5).

### Supplementary Figures

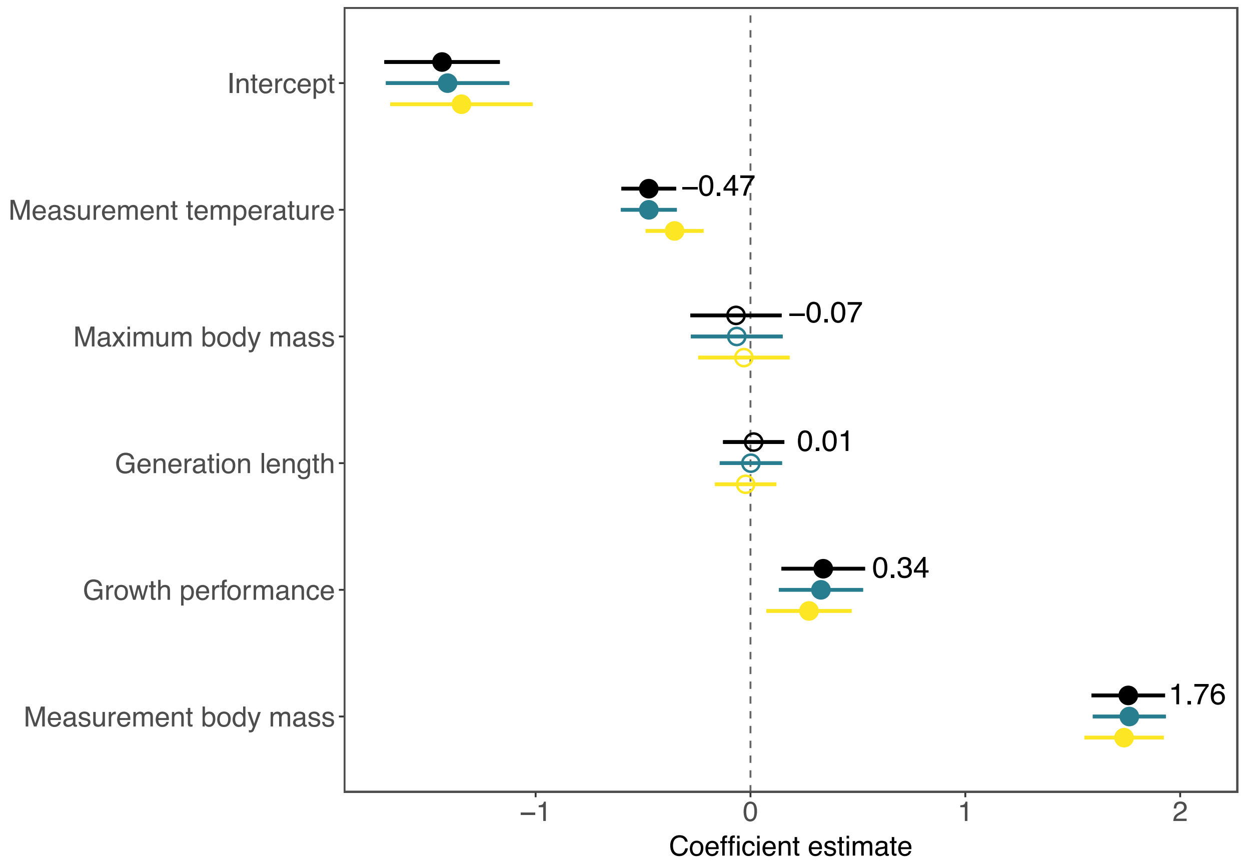

**Figure S1.** Coefficient plot of the relative effects of measurement body mass, measurement temperature, growth performance, maximum body mass, and generation length on RMR from global PGLS models (n = 104). Models were run using RMR data from three datasets – either the sample size dataset (black), the mass dataset (green), or the temperature dataset (yellow). Values next to the black points are the resulting model coefficients from the sample size dataset. Measurement temperature was modeled as inverse temperature, all variables except measurement temperature were $log_{10}$-transformed, and all explanatory variables were standardized. Horizontal lines represent 95% confidence intervals.

### Supplementary Tables

**Table S1.** Comparisons of phylogenetic generalized least squares models investigating how life history traits explain variation in resting metabolic rate (RMR) across fishes, while accounting for measurement body mass ($M_b$) and measurement temperature ($T$). Life history traits are maximum body mass ($M_{max}$), generation length ($GL$), and growth performance ($\phi$’). All variables were $log_{10}$-transformed, and measurement temperature was modeled as standardized inverse temperature. RMR data was from either the sample size dataset (1), the mass dataset (2), or the temperature dataset (3).

| | **Model:** RMR ~ | **$\lambda$** | **df** | **AICc** | **ΔAICc** | **$w_i$** |
|---|---|---|---|---|---|---|
| *1) Sample size* dataset | | | | | | |
| | $M_b + T + \phi$’ | 0.56 | 4 | 16.84 | 0.00 | 0.99 |
| | $M_b + T$ | 0.71 | 3 | 27.47 | 10.62 | 0.00 |
| | $M_b + T + M_{max}$ | 0.69 | 4 | 27.80 | 10.96 | 0.00 |
| | $M_b + T + GL$ | 0.72 | 4 | 29.58 | 12.74 | 0.00 |
| *2) Mass dataset* | | | | | | |
| | $M_b + T + \phi$’ | 0.61 | 4 | 19.35 | 0.00 | 0.99 |
| | $M_b + T$ | 0.71 | 3 | 29.49 | 10.14 | 0.01 |

| | | | | | | |
|---|---|---|---|---|---|---|
| | $M_b + T + M_{max}$ | 0.69 | 4 | 29.93 | 10.58 | 0.00 |
| | $M_b + T + GL$ | 0.71 | 4 | 31.65 | 12.30 | 0.00 |
| *3) Temperature dataset* | | | | | | |
| | $M_b + T + \phi'$ | 0.71 | 4 | 17.62 | 0.00 | 0.90 |
| | $M_b + T$ | 0.82 | 3 | 23.13 | 5.51 | 0.06 |
| | $M_b + T + M_{max}$ | 0.81 | 4 | 24.80 | 7.18 | 0.02 |
| | $M_b + T + GL$ | 0.81 | 4 | 25.05 | 7.43 | 0.02 |

**Table S2.** Coefficients and 95% confidence intervals (CIs) from phylogenetic generalized least squares models investigating how life history traits explain variation in resting metabolic rate (RMR) across fishes (n = 104), while accounting for measurement body mass and measurement temperature. The 'null' model included only measurement body mass and measurement temperature as explanatory variables. Life history traits are maximum body mass ($M_{max}$), generation length ($GL$), growth performance ($\phi$'), age at maturity ($T_{mat}$), maximum age ($T_{max}$), growth rate ($k$), and asymptotic length ($L_\infty$). RMR data used to estimate these coefficients came from the sample size dataset. All variables were $\log_{10}$-transformed, and measurement temperature was modeled as standardized inverse temperature.

| | **Intercept** | | | **Measurement body mass** | | | **Measurement temperature** | | | **Life history trait** | | |
|---|---|---|---|---|---|---|---|---|---|---|---|---|
| | | **95% CI** | | | **95% CI** | | | **95% CI** | | | **95% CI** | |
| | | **lower** | **upper** | | **lower** | **upper** | | **lower** | **upper** | | **lower** | **upper** |
| null | -3.26 | -3.63 | -2.89 | 0.87 | 0.81 | 0.92 | -0.52 | -0.65 | -0.39 | – | – | – |
| $M_{max}$ | -3.33 | -3.70 | -2.95 | 0.83 | 0.75 | 0.91 | -0.52 | -0.65 | -0.39 | 0.04 | -0.02 | 0.11 |
| $GL$ | -3.28 | -3.72 | -2.85 | 0.87 | 0.81 | 0.92 | -0.52 | -0.66 | -0.39 | 0.03 | -0.21 | 0.26 |
| $\phi'$ | -3.76 | -4.15 | -3.38 | 0.79 | 0.72 | 0.86 | -0.48 | -0.60 | -0.36 | 0.24 | 0.12 | 0.36 |
| $T_{mat}$ | -3.24 | -3.62 | -2.87 | 0.87 | 0.81 | 0.94 | -0.51 | -0.65 | -0.36 | -0.04 | -0.22 | 0.15 |
| $T_{max}$ | -3.30 | -3.73 | -2.87 | 0.86 | 0.81 | 0.92 | -0.53 | -0.66 | -0.39 | 0.04 | -0.16 | 0.25 |
| $k$ | -3.19 | -3.56 | -2.82 | 0.88 | 0.82 | 0.93 | -0.49 | -0.62 | -0.35 | 0.12 | -0.03 | 0.28 |
| $L_\infty$ | -3.49 | -3.94 | -3.03 | 0.83 | 0.76 | 0.91 | -0.53 | -0.66 | -0.40 | 0.17 | -0.05 | 0.38 |

**Table S3.** Coefficients and 95% confidence intervals from phylogenetic generalized least squares models investigating how life history traits explain variation in resting metabolic rate (RMR) across fishes (n = 104), while accounting for measurement body mass ($M_b$) and measurement temperature ($T$). Life history traits are maximum body mass ($M_{max}$), generation length ($GL$), and growth performance ($\phi$'). RMR data used to estimate these coefficients came from either the mass dataset or the temperature dataset. All variables were $log_{10}$-transformed, and measurement temperature was modeled as standardized inverse temperature.

| | **Mass dataset** | | | **Temperature dataset** | | |
|---|---|---|---|---|---|---|
| | | **95% CI** | | | **95% CI** | |
| | | **lower** | **upper** | | **lower** | **upper** |
| $RMR \sim M_b + T$ | | | | | | |
| Intercept | -3.26 | -3.64 | -2.89 | -3.25 | -3.67 | -2.82 |
| $M_b$ | 0.87 | 0.82 | 0.93 | 0.88 | 0.82 | 0.93 |
| $T$ | -0.52 | -0.65 | -0.39 | -0.41 | -0.55 | -0.28 |
| $RMR \sim M_b + T + M_{max}$ | | | | | | |
| Intercept | -3.33 | -3.70 | -2.95 | -3.28 | -3.71 | -2.85 |
| $M_b$ | 0.84 | 0.76 | 0.92 | 0.86 | 0.77 | 0.94 |
| $T$ | -0.52 | -0.65 | -0.39 | -0.41 | -0.55 | -0.27 |
| $M_{max}$ | 0.04 | -0.02 | 0.11 | 0.02 | -0.04 | 0.09 |
| $RMR \sim M_b + T + GL$ | | | | | | |
| Intercept | -3.26 | -3.70 | -2.83 | -3.19 | -3.67 | -2.72 |
| $M_b$ | 0.87 | 0.81 | 0.93 | 0.88 | 0.82 | 0.94 |
| $T$ | -0.52 | -0.66 | -0.38 | -0.41 | -0.55 | -0.27 |
| $GL$ | 0.00 | -0.24 | 0.24 | -0.06 | -0.29 | 0.18 |
| $RMR \sim M_b + T + \phi'$ | | | | | | |
| Intercept | -3.75 | -4.15 | -3.35 | -3.66 | -4.09 | -3.22 |
| $M_b$ | 0.80 | 0.73 | 0.87 | 0.80 | 0.73 | 0.88 |
| $T$ | -0.48 | -0.60 | -0.36 | -0.36 | -0.49 | -0.23 |
| $\phi'$ | 0.23 | 0.11 | 0.36 | 0.20 | 0.07 | 0.34 |

**Table S4.** Coefficients and 95% confidence intervals from phylogenetic generalized least squares models investigating whether weight-based growth performance ($\phi$) explains variation in resting metabolic rate (RMR), while accounting for measurement body mass ($M_b$) and measurement temperature ($T$). Growth performance was calculated using (A) group-and species-specific length-weight regressions as well as (B) species-specific, but not group-specific, length-weight regression equations. Measurement temperature was modeled as standardized inverse temperature and all variables were $log_{10}$-transformed.

| | **A (n = 44)** | | | **B (n = 84)** | | |
|---|---|---|---|---|---|---|
| | | **95% CI** | | | **95% CI** | |
| | | **lower** | **upper** | | **lower** | **upper** |
| Intercept | -3.28 | -3.69 | -2.88 | -3.35 | -3.67 | -3.02 |
| $M_b$ | 0.82 | 0.71 | 0.94 | 0.82 | 0.75 | 0.89 |
| $T$ | -0.45 | -0.61 | -0.28 | -0.50 | -0.63 | -0.38 |
| $\phi$ | 0.09 | -0.09 | 0.27 | 0.13 | 0.01 | 0.25 |

**Table S5.** Comparisons of phylogenetic generalized least squares models investigating how life history traits explain variation in resting metabolic rate (RMR) across fishes, while accounting for measurement body mass ($M_b$) and measurement temperature ($T$). Life history traits are maximum body mass ($M_{max}$), generation length ($GL$), length-based growth performance ($\phi$'), and weight-based growth performance ($\phi$). All variables were $log_{10}$-transformed, and measurement temperature was modeled as standardized inverse temperature. Weight-based growth performance was calculated using (A) group- and species-specific length-weight regression equations as well as (B) species-specific, but not group-specific, length-weight regression equations.

| | **Model:** RMR ~ | $\lambda$ | df | AICc | ΔAICc | $w_i$ |
|---|---|---|---|---|---|---|
| A (n = 44) | | | | | | |
| | $M_b + T$ | 0.81 | 3 | 5.33 | 0.00 | 0.37 |
| | $M_b + T + \phi'$ | 0.82 | 4 | 6.39 | 1.05 | 0.22 |
| | $M_b + T + \phi$ | 0.71 | 4 | 6.77 | 1.44 | 0.18 |
| | $M_b + T + M_{max}$ | 0.83 | 4 | 7.55 | 2.21 | 0.12 |
| | $M_b + T + GL$ | 0.82 | 4 | 7.74 | 2.41 | 0.11 |
| B (n = 84) | | | | | | |
| | $M_b + T + \phi'$ | 0.74 | 4 | -2.38 | 0.00 | 0.42 |
| | $M_b + T + \phi$ | 0.71 | 4 | -2.05 | 0.33 | 0.35 |
| | $M_b + T$ | 0.77 | 3 | 0.01 | 2.39 | 0.13 |
| | $M_b + T + M_{max}$ | 0.77 | 4 | 1.75 | 4.13 | 0.05 |
| | $M_b + T + GL$ | 0.78 | 4 | 2.03 | 4.41 | 0.05 |

**Table S6.** Coefficients and AICc comparisons for phylogenetic generalized least squares models investigating how life history traits explain variation in resting metabolic rate (RMR) while excluding life history data from proxy species (n = 76) and while accounting for measurement body mass ($M_b$) and measurement temperature ($T$). Life history traits are growth performance ($\phi'$), maximum body mass ($M_{max}$), and generation length ($GL$). RMR data for these analyses came from the sample size dataset. All variables were $\log_{10}$-transformed, and measurement temperature was modeled as standardized inverse temperature.

| | **Estimate** | **lower 95% CI** | **upper 95% CI** | $\lambda$ | **df** | **AICc** | **ΔAICc** | $w_i$ |
|---|---|---|---|---|---|---|---|---|
| $RMR \sim M_b + T + \phi'$ | | | | 0.72 | 4 | 1.40 | 0.00 | 0.92 |
| Intercept | -3.61 | -4.04 | -3.17 | | | | | |
| $M_b$ | 0.78 | 0.71 | 0.86 | | | | | |
| $T$ | -0.50 | -0.64 | -0.36 | | | | | |
| $\phi'$ | 0.19 | 0.07 | 0.32 | | | | | |
| $RMR \sim M_b + T$ | | | | 0.79 | 3 | 7.95 | 6.55 | 0.03 |
| Intercept | -3.18 | -3.59 | -2.78 | | | | | |

| | | | | | | | | |
|---|---|---|---|---|---|---|---|---|
| $M_b$ | 0.84 | 0.78 | 0.91 | | | | | |
| $T$ | -0.55 | -0.70 | -0.41 | | | | | |
| $RMR \sim M_b + T + M_{max}$ | | | | 0.79 | 4 | 8.22 | 6.83 | 0.03 |
| Intercept | -3.26 | -3.67 | -2.85 | | | | | |
| $M_b$ | 0.81 | 0.73 | 0.89 | | | | | |
| $T$ | -0.55 | -0.70 | -0.41 | | | | | |
| $M_{max}$ | 0.04 | -0.02 | 0.11 | | | | | |
| $RMR \sim M_b + T + GL$ | | | | 0.80 | 4 | 10.15 | 8.75 | 0.01 |
| Intercept | -3.20 | -3.67 | -2.74 | | | | | |
| $M_b$ | 0.84 | 0.78 | 0.91 | | | | | |
| $T$ | -0.56 | -0.71 | -0.41 | | | | | |
| $GL$ | 0.02 | -0.22 | 0.26 | | | | | |
